# Multiplex CRISPRi-Cas9 silencing of planktonic and stage-specific biofilm genes in *Enterococcus faecalis*

**DOI:** 10.1101/2020.04.30.071571

**Authors:** Irina Afonina, June Ong, Jerome Chua, Timothy Lu, Kimberly A. Kline

**Affiliations:** Singapore–MIT Alliance for Research and Technology, Antimicrobial Drug Resistance Interdisciplinary Research Group, Singapore 138602; School of Biological Sciences, Nanyang Technological University, 60 Nanyang Drive, Singapore 637551; Electrical Engineering and Computer Science, MIT, Cambridge, MA 02139, USA; Department of Biological Engineering, MIT, Cambridge, MA 02139, USA.; Singapore Centre for Environmental Life Science Engineering, Nanyang Technological University, 60 Nanyang Drive, Singapore 637551

**Keywords:** *Enterococcus faecalis*, CRISPR interference, biofilms, gene essentiality, Ebp pili

## Abstract

*Enterococcus faecalis* is an opportunistic pathogen, which can cause multidrug-resistant life-threatening infections. Gaining a complete understanding of enterococcal pathogenesis is a crucial step in identifying a strategy to effectively treat enterococcal infections. However, bacterial pathogenesis is a complex process often involving a combination of genes and multi-level regulation. Compared to established knockout methodologies, CRISPRi approaches enable rapid and efficient silencing of genes to interrogate gene products and pathways involved in pathogenesis. As opposed to traditional gene inactivation approaches, CRISPRi can also be quickly repurposed for multiplexing or used to study essential genes. Here we have developed a novel dual-vector nisin-inducible CRISPRi system in *E. faecalis* that can efficiently silence via both non-template and template strand targeting. Since nisin-controlled gene expression system is functional in various Gram-positive bacteria, the developed CRISPRi tool can be extended to other genera. This system can be applied to study essential genes, genes involved in antimicrobial resistance, and genes involved in biofilm formation and persistence. The system is robust, and can be scaled up for high-throughput screens or combinatorial targeting. This tool substantially enhances our ability to study enterococcal biology and pathogenesis, host-bacteria interactions, and inter-species communication.

**IMPORTANCE:** *Enterococcus faecalis* causes multidrug resistant life-threatening infections, and is often co-isolated with other pathogenic bacteria from polymicrobial biofilm-associated infections. Genetic tools to dissect complex interactions in mixed microbial communities are largely limited to transposon mutagenesis and traditional time- and labour-intensive allelic exchange methods. Built upon streptococcal dCas9, we developed an easily-modifiable, inducible CRISPRi system for *E. faecalis* that can efficiently silence single and multiple genes. This system can silence genes involved in biofilm formation, antibiotic resistance, and can be used to interrogate gene essentiality. Uniquely, this tool is optimized to study genes important for biofilm initiation, maturation, and maintenance, and can be used to perturb pre-formed biofilms. This system will be valuable to rapidly and efficiently investigate a wide range of aspects of complex enterococcal biology.

## INTRODUCTION

Enterococci are Gram-positive, opportunistic pathogens that are the second leading cause of the hospital-acquired infections (HAI) [1]. Within the *Enterococcus* species, *Enterococcus faecalis* and *Enterococcus faecium* are most commonly isolated from human infection, and *E. faecalis* is most frequently isolated in HAI [2]. *E. faecalis* causes life-threatening endocarditis, bacteraemia, wound infection, and medical device-associated infections including catheter-associated urinary tract infections [3, 4]. Many of these infections are biofilm-associated, resulting in their increased tolerance to antibiotic clearance. In addition, enterococci are intrinsically resistant to multiple classes of antibiotics and rapidly acquire resistance through mutation and horizontal gene transfer, further rendering these infections difficult to treat [2, 5]. Understanding the mechanisms of biofilm formation, antimicrobial resistance, host immune evasion, and inter-species communication is crucial to more effectively manage and treat enterococcal infections. However, biofilms and antimicrobial resistance involve complex gene pathways comprised of multiple genes, which makes it difficult to study with current available tool that designed to study one gene at a time. To date, genetic tools to study enterococcal biology are limited to transposon mutagenesis and allelic-exchange gene inactivation or deletion, both of which are laborious, time-consuming, and only scalable in a decelerated step-by-step manner [6–8].

Clustered, regularly interspaced, short palindromic repeat (CRISPR) loci coupled with CRISPR-associated (Cas) proteins were first described to confer bacterial adaptive immunity against bacteriophages and invading plasmids [9–11]. Since repurposing of CRISPR-Cas systems for gene editing, the toolbox for genetic manipulation in bacteria is expanding [12, 13]. The well-studied Type II CRISPR-Cas system consists of a DNA endonuclease (Cas9) that is guided to the bacterial chromosome by a short 20 nt single guide RNA (sgRNA), where they generate a double-stranded DNA break by recognizing a 2–6-base pair DNA sequence called a protospacer-adjacent motif (PAM) that immediately follows the targeted gene [14]. Lack of an efficient mechanism for non-homologous end joining in bacteria makes CRISPR-Cas9 lethal, which inspired the repurposing of CRISPR-Cas for antimicrobial therapy [15–18]. CRISPR interference (CRISPRi) takes advantage of a catalytically inactive or “dead” Cas9 (dCas9) that sterically blocks transcription elongation to control gene expression [19, 20]. CRISPRi also enables large-scale genome-wide studies and the simultaneous silencing of multiple genes, and has been successfully implemented in *Escherichia coli, Bacillus subtilis*, and *Streptococcus pneumonia* where high-throughput screens identified essential bacterial genes [20–22]. The well-characterized dCas9 from *Streptococcus pyogenes* is generally used for genetic perturbation studies because the Cas9 handle, a 42 nt Cas9-binding hairpin, and PAM sequence are well defined [13, 19, 23]. However, because *S. pyogenes* dCas9 performance varies in different species, with low knockdown efficiency and proteotoxicity in *Mycobacteria tuberculosis* for example, dCas9 from other species such as *Streptococcus thermophilus* have also been effectively used for CRISPRi [24, 25].

*E. faecalis* encodes a Type II CRISPR-Cas9 system with a canonical PAM of NGG (where N indicates any nucleotide) [26]. In *E. faecalis*, basal levels of chromosomally-encoded Cas9 guided to an incoming plasmid via a specific CRISPR RNA and transactivating RNA (cr-RNA-tracrRNA) complex is insufficient to fully prevent the conjugation of a foreign conjugative plasmid [27]. However, native chromosomally encoded CRISPR-Cas9 also has been successfully used to target antimicrobial resistance genes *in vivo* with a significant reduction of the targeted population compared to a ∆*cas9* control [18, 26]. Overexpression of Cas9 significantly improves enterococcal immune capacity by diminishing the rates of plasmid transfer to non-detectable levels *in vitro* [27].

While native chromosomal CRISPR-Cas9 has been used for targeted mutagenesis in *E. faecalis*, a CRISPRi tool for a high-throughput scalable genetic control studies is still lacking. Here we developed a dual-vector nisin-inducible system for *E. faecalis* that can efficiently silence single genes and whole operons. This system can also be easily multiplexed to repress multiple genes at the same time. We show that the CRISPRi system can be used to study the genes involved in biofilm formation, antimicrobial resistance, as well as essential genes. Importantly, we report effective selective CRISPRi silencing by targeting either the non-template or template-DNA strand, expanding our understanding for how CRISPRi can work. In addition, we demonstrate the reduction of pre-formed biofilms through CRISPRi targeting of biofilm-associated genes. Simultaneous silencing of multiple genes and essential genes provides an easily engineered tool to dissect mechanisms of enterococcal pathogenesis, antimicrobial resistance, host-pathogen interactions, and cross-species communication.

## MATERIALS AND METHODS

### Bacterial strains and media conditions

The strains and plasmids used in the study are listed in Table 1. *E. faecalis* strains were grown statically at 37°C in tryptone soy broth (Oxoid, UK) supplemented with 10 mM glucose (TSBG) for biofilm studies, Mueller Hinton broth 2 (MHC-2, Merck, USA) for bacitracin susceptibility tests, and in brain heart infusion media (BHI; Merck, USA) or BHI agar (Merck, USA) for the rest of the experiments. *E. coli* was grown in Luria-Bertani Broth Miller (LB; BD, Difco, USA) at 37°C, 200 rpm shaking. Erythromycin (100 µg/ml) was used to maintain pMSP3545 plasmid in *E. faecalis*; kanamycin (500 µg/ml for *E. faecalis* and 50 µg/ml for *E. coli*) was used to maintained pGCP123 and its derivatives. Nisin (Sigma, USA) stock solution was prepared as 0.1 mg/ml, by dilution in deionized water. The nisin solution was then filter sterilized through a 0.22 µm filter, aliquoted, and frozen at −20°C. When needed an aliquot was thawed and used once.

**Table 1.** Bacterial strains used in the study.

| Bacterial strains | Description | Reference |
| --- | --- | --- |
| OG1RF | Human oral isolate OG1 | [60] |
| SD234 | GFP-tagged OG1RF | [37] |
| SD234::dCas9 | Catalytically inactive dCas9; mutations D10A, H840A | This study |
| $\Delta ebpABC$ | Contains in-frame-deletion of *ebpABC* operon | [42] |
| *croR*::Tn | Transposon mutant of *croR* | [61] |
| $\Delta srtA$ | Contains in-frame-deletion of *srtA* gene | [30] |
| **Plasmids** |  |  |
| pGCP123 | Small shuttle vector for sgRNA expression | [8] |
| pGCP213 | Temperature-sensitive integration vector for allelic exchange | [8] |
| pMSP3545 | Encodes nisin two-component system and nisin inducible promoter *nisA* | a gift from Gary Dunny (Addgene plasmid # 46888) [32] |
| pMSP3545-dCas9<sub>Str</sub> | Encodes dCas9<sub>Str</sub> under nisin inducible promoter *nisA* | this study |
| pdCas9-bacteria | Catalytically inactive Cas9 from *Streptococcus pyogenes* | a gift from Stanley Qi (Addgene plasmid # 44249) [23] |

### Genetic manipulations

To abolish endonuclease activity of native enterococcal Cas9_EF_, we first aligned *csn1* (OG1RF_10404) to *S. pyogenes* Cas9 (BlastP, NCBI) and identified conserved catalytic D10 and H852 residues within the RuvC1 active site and HNH endonuclease domains, respectively [13]. Both of the amino acids are essential for cleaving the template and non-template DNA strands [13]. To create D10A and H852A substitutions and introduce 1 kb flanking region to Cas9_EF_ for the subsequent allelic exchange, we amplified 3 fragments from the bacterial chromosome with primers pairs 1/2, 3/4 and 5/6 listed in Table 2 and performed splice overlap extension (SOE) PCR of the 3 fragments with 1/6 primer pair. The resulting product was introduced into the PstI/KpnI digested temperature-sensitive vector pGCP213 by In-Fusion (Takaro Bio, Japan). The resulting pGCP213-dCas9EF vector was verified by sequencing and used for allelic exchange to generate SD234::dCas9 as described previously [8]. The presence of both mutations D10A and H852A in the *csn1* gene encoding Cas9_EF_ were verified by PCR and sequencing with primer pairs 7/8 and 9/10.

**Table 2.**
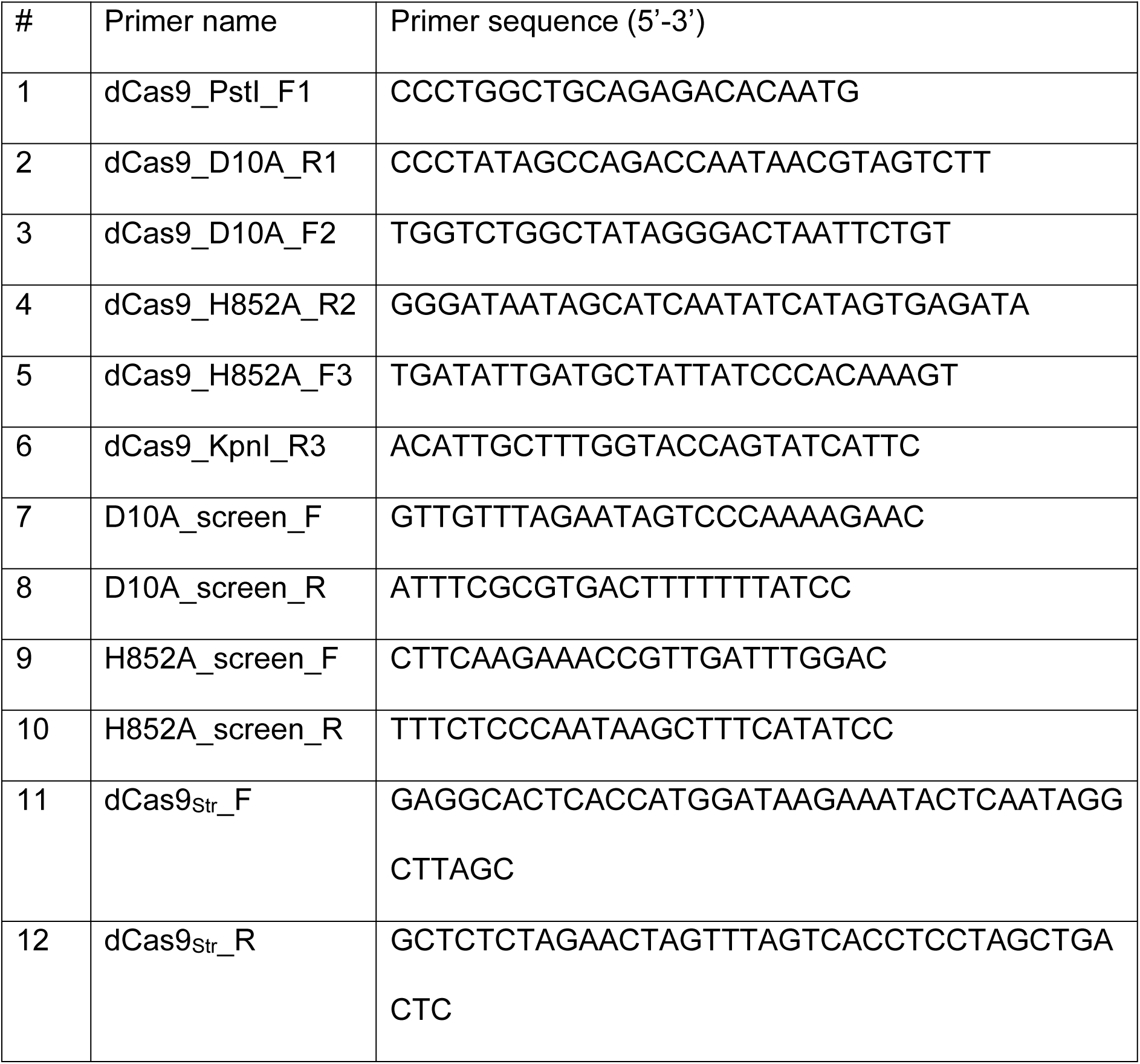
Primers used in the study.

To generate pMSP3545-dCas9_Str_, dCas9_Str_ was amplified from pdCas9-bacteria (Addgene #44249) using the primer pair 11/12, and the purified product was introduced by In-Fusion into pMSP3545 (Addgene #46888) digested with SpeI and NcoI. To generate sgRNA expression vectors, a 307 bp gBlock (IDT, USA) consisting of the *nisA* promoter linked directly to a 20 nt sgRNA linked to the dCas9 scaffold (**Figure S1**) was introduced by In-Fusion into pGCP123 digested with BglII and NotI. The gBlock contains 4 restriction sites (BglII, BamHI, EcoRI and MfeI) for the generation of a 2-wise library through restriction-ligation reactions, and an 8 nt unique barcode for high-throughput screens coupled with amplicon sequencing (**Figure S1**) [28]. sgRNA sequences were selected using the CHOPCHOP database with a zero off-target score and minimal self-complementarity (0-1) (Table 3) [29].

**Table 3.**
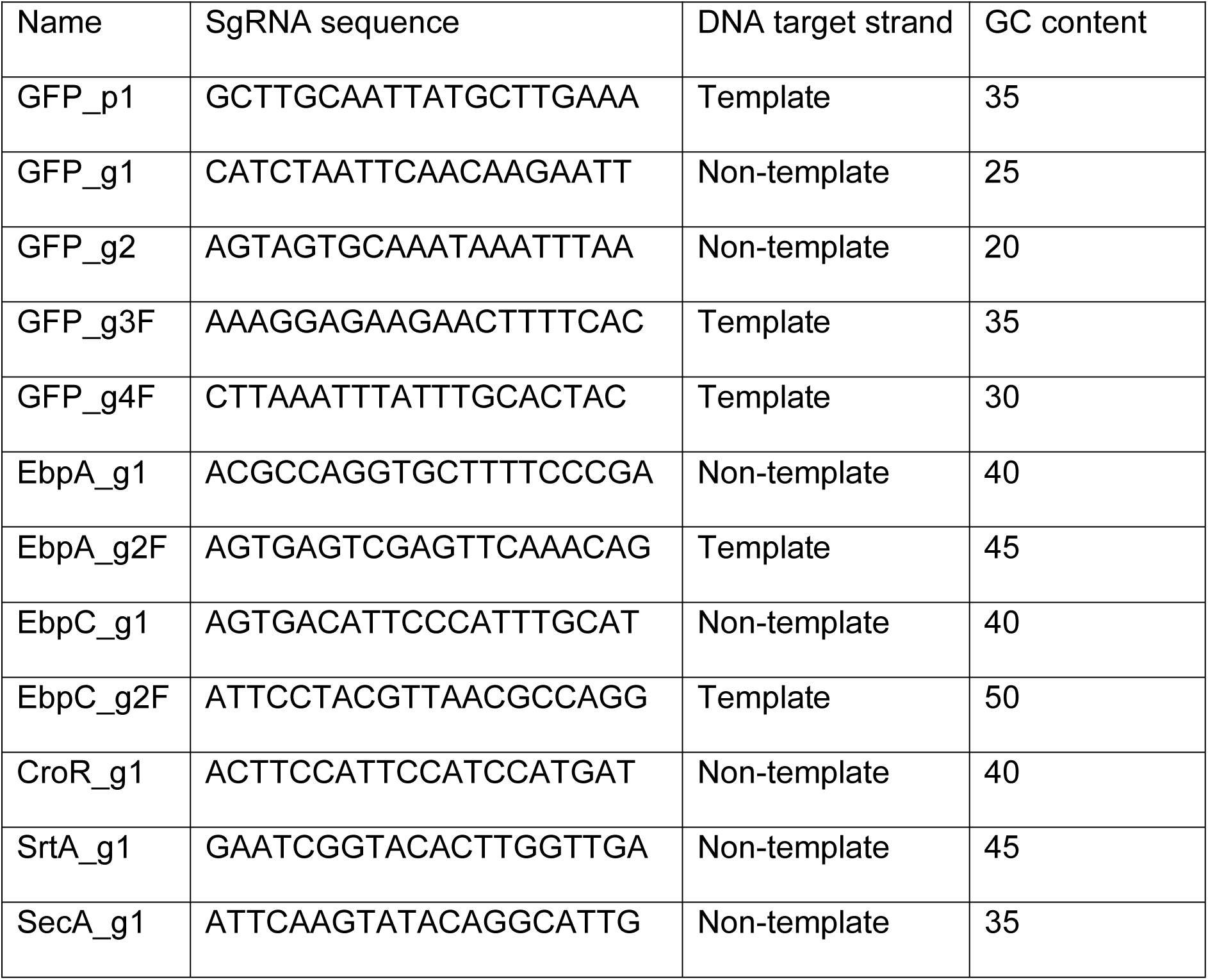
SgRNAs used in the study

### Flow cytometry

A single colony was inoculated and grown overnight with or without the addition of nisin. The following day cultures were diluted 1:30 in 1,160 µl of fresh media in a 2 ml tube and grown for 3 hours statically at 37°C. After incubation, the cells were collected by centrifugation, resuspended in 1 ml PBS, and analysed on an Attune NxT Flow Cytometer. The percentage of GFP expressing cells was determined by proprietary Attune NxT flow cytometry software from 500 000 events based on polygonal gating on EfdCas9 empty vector control.

### Western blot

A single colony was selected from agar plates, inoculated into liquid media, and grown overnight with or without nisin induction. Overnight cultures were then diluted 1:10 in fresh media in the presence of antibiotics and nisin where appropriate and grown until mid-log phase. Samples were normalized to OD_600nm_ 0.6, pelleted by centrifugation, and the pellet was resuspended in 75 µl of 10 mg/ml of lysozyme in lysozyme buffer (10 mM Tris-HCl pH 8, 50 mM NaCl, 1 mM EDTA, 0.75 M sucrose) and incubated at 37°C for 1 hour. After lysozyme treatment, 25 µL of 4X NuPAGE^®^ LDS sample buffer (Invitrogen, USA) was added to the samples, the samples were heated at 95°C for 10 min, and stored at −20°C until analysis.

For immunoblot analysis, 10 µl of each sample was loaded onto a 4-12% gradient NuPAGE^®^ Bis-Tris mini gel for SecA or SrtA, or a 3-8% gradient NuPAGE^®^ Bis-Tris mini gel for EbpA, and run in 1xMOPS (for 4-12% gel) or tris-acetate (for 3-8% gel) SDS running buffer, respectively, in a XCell*SureLock^®^*Mini-Cell for 50 min at 200V. Proteins from the gel were then transferred to a membrane using the iBlot^TM^ Dry Blotting system. The membrane was then blocked with 3% bovine serum albumin (BSA) in phosphate buffer saline with 0.05% Tween20 (PBS-T) for one hour, with shaking at RT. SecA, SrtA and EbpA were detected using custom antibodies raised in rabbit, mouse and guinea pig respectively [8, 30]. Cas9 was detected using a monoclonal mouse anti-Cas9 antibody (Abcam). Appropriate IgG secondary horseradish peroxidase (HRP)-conjugated antibodies (all Thermo Scientific, Singapore) were used for detection.

### Biofilm assay

Overnight bacterial cultures were washed and normalized to OD_600nm_ 0.7 as described previously [31]. 5000 CFU/well were inoculated in TSBG in a 96-well flat-bottom transparent microtiter plate (Thermo Scientific, Waltman, MA, USA), and incubated at 37°C under static conditions for 24 hours. After removal of planktonic cells, the adherent biofilm biomass was stained using 0.1% w/v crystal violet (Sigma-Aldrich, St Louis, MO, USA) at 4°C for 30 minutes. The microtiter plate was washed twice with PBS followed by crystal violet solubilization with ethanol: acetone (4:1) for 45 minutes at room temperature. Quantification of adherent biofilm biomass was measured by absorbance at OD_595nm_ using a Tecan Infinite 200 PRO spectrophotometer (Tecan Group Ltd., Männedorf, Switzerland).

### Bacitracin susceptibility determination

The minimal inhibitory concentration (MIC) of bacitracin was determined in liquid MHB II media in a 96-well plate. Two-fold serial dilutions from 128 µg/ml to 4 µg/ml of bacitracin were prepared in triplicate from a 512 µg/ml bacitracin stock. Overnight cultures of bacteria were normalized to OD600_nm_ 0.7 and 8 µl of inoculated in each well containing 200 µl TSBG media, with the final concentration of 10^5^ CFU/ml. Plates were incubated at 37°C for 16 hours. The next day the MIC was determined by visually assessing turbidity. The lowest concentration of the antibiotic that prevented growth was recorded as the MIC.

### Growth curve assessment

Overnight cultures were washed in PBS and normalized to OD_600nm_ 0.7. Normalized cultures were inoculated into 200 ul BHI media at a ratio 1:25. Three biological replicates and 4 technical replicates were performed for each culture. The 96-well plates were incubated at 37°C for 16 hours using the BioTek synergy 4 (BioTek, USA) plate reader. Optical density was taken at OD_600nm_ at 30 min intervals to determine the growth curve of each culture.

### Statistical Analysis

Statistical analyses were performed using GraphPad Prism software (Version 6.05 for Windows, California, United States). All experiments were performed at least in three biological replicates and the mean value was calculated. All graphs show the standard deviation from independent experiments. Statistical analysis was performed by the unpaired t-test using GraphPad (* p<0.05, ** p<0.01, *** p<0.001; **** p<0.0001, ns: p>0.05). *P*-values less than 0.05 were deemed significant.

## RESULTS

### Construction of a dual-vector CRISPRi system in *E. faecalis*

To design a scalable CRISPRi expression system, we used the catalytically inactive *S. pyogenes* Cas9 (dCas9) with its well-defined Cas9 scaffold sequence. We cloned *dcas9* from pdCas9-bacteria (Addgene) under the nisin inducible promoter *nisA* in pMSP3545, which also encodes the nisin responsive NisKR two-component system, to generate pMSP3545-dCas9 [23, 32] (Figure 1A). Barcoded gRNA sequences with a dCas9 handle under control of the same *nisA* promoter were synthesized as gBlocks (IDT, USA) and cloned into the pGCP123 expression vector by InFusion reaction to generate pGCP123-sgRNA [8, 23, 33]. Both plasmids were transformed into *E. faecalis*. Upon addition of nisin to the media, NisK is activated and phosphorylates NisR, which binds to the *nisA* promoter to drive expression of dCas9 from pMSP3545-dCas9 and sgRNA from pGCP123-sgRNA (Figure 1A) [33]. The strength of the *nisA* promoter is dose-dependent and peaks at 25 ng/ml of nisin (Figure 1B).

**Figure 1.**
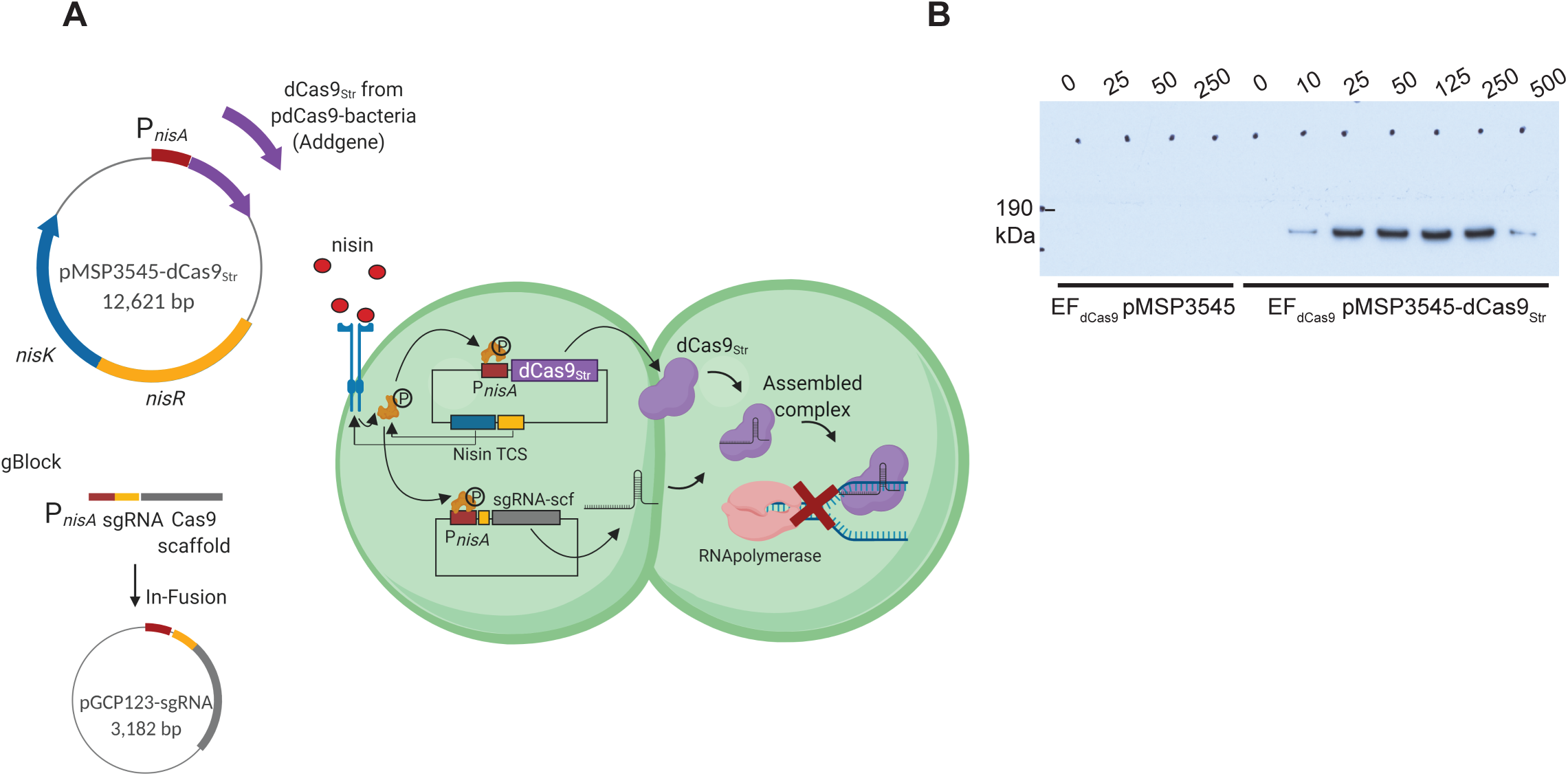
Nisin-inducible dual-vector CRISPRi in *Enterococcus faecalis*. **A.** Schematic diagram of CRISPRi system in *Enterococcus faecalis.* A two-plasmid system consisting of a small (3,182 kb) vector, pGCP123, for sgRNA expression and a 12,621kb plasmid, pMSP3545-dCas9_Str_, for dCas9_Str_ expression. The sgRNA and dCas9_Str_ are expressed from a nisin-inducible promoter *nisA* that is activated upon addition of nisin to the media through the NisKR two-component system encoded on pMSP3545-dCas9_Str_. The assembled sgRNA-dCas9 complex blocks gene transcription by binding to DNA and blocking RNA polymerase. Image created with BioRender.com. **B.** Western blot with anti-Cas9_Str_ antibody on induced EF_dCas9_ pMSP3545 (empty vector) and EF_dCas9_ pMSP3545-dCas9_Str_ at nisin concentration 0-500 ng/ml.

CRISPR-Cas systems are categorized into six major types (I through VI), with each having a type-specific *cas* gene [34]. The CRISPR-Cas system in *E. faecalis* is a Type II system, which possesses the type-specific gene *cas9*, that in OG1RF is encoded by *csn1* (OG1RF_10404) [35]. Since streptococcal Cas9_Str_ is orthologous to enterococcal Cas9 (NCBI, Blastp) and shares the same PAM – NGG [26, 36], we first tested whether the chromosomally-encoded, inactivate *E. faecalis* Cas9 can recognize the streptococcal dCas9 handle and interfere with the episomal inducible CRISPRi system. To test this, we compared CRISPRi activity of streptococcal dCas9_Str_ to inactivated native enterococcal dCas9_EF_. In *E. faecalis* SD234 (strain OG1RF expressing *gfp* on the chromosome [37]) we mutated catalytic residues D10A and H852A to generate the strain SD234::dCas9 (EF*_dCas9_*). The two catalytic residues correspond to streptococcal catalytic residues D10 and H840 in the RuvC-I domain and HNH domain, respectively, that are responsible for non-template and template DNA strand cleavage [13]. We then co-transformed GFP_g2, encoding a sgRNA that targets the chromosomally encoded *gfp*, together with either the pMSP3545 empty vector control or with pMSP3545-dCas9_Str_, into EF_dCas9_. We monitored GFP signal using flow cytometry, and upon nisin induction, we observed only 1% of cells transformed with p3545-dCas9_Str_ remained GFP-positive compared to 98% GFP positive cells in the empty vector control (**Figure S2**). Since inactivated dCas9 from *E. faecalis* does not recognize the streptococcal scaffold to silence *gfp* in the empty vector control, nor does it interfere with *gfp* silencing by dCas9Str, then we reason that the native (catalytically active) enterococcal Cas9 would also not bind to the scaffold, since only catalytic and not binding residues are mutated. These results demonstrate that dCas9_EF_ from *E. faecalis* does not interfere with scaffold recognition, and subsequent gene silencing, by streptococcal dCas9.

### CRISPRi silencing of chromosomal gene via template or non-template strand targeting

We next tested two parameters that can be potentially optimized for efficient CRISPRi targeting, namely GC content of the sgRNA and guide position within the gene and its promoter region [23, 24, 38]. We also tested template and nontemplate DNA strand targeting to determine whether only nontemplate strand targeting is efficient in *E. faecalis*, as has been shown in *E. coli*, *Pseudomonas aeruginosa, Mycobacterium tuberculosis, and Caulobacter crescentus* [23–25, 39]. We designed guides to test the ability of CRISPRi to silence *gfp* gene by targeting a) a promoter region with GFP_p1 (35% GC); b) a non-template DNA strand with GFP_g1 (25% GC) and GFP_g2 (20% GC); c) a template DNA strand with GFP_g3F (35% GC) and GFP_g4F (30% GC) (Figure 2A). To determine the expression conditions for maximal targeting efficiency, we used a saturating concentration of nisin of 50 ng/ml and compared planktonic bacteria subcultured without (-) or with (+) nisin for 2 hours. We observed only partial silencing (up to 70%) for 4 of the 5 guides when bacteria were induced for 2 hours (Figure 2B). To improve silencing, we pre-sensitized bacteria with nisin induction overnight before subculturing the bacteria into fresh media again with nisin for 2 hours (++). Pre-sensitizing the bacteria universally increased the silencing efficiency for all active guides from 70% to 99% (Figure 2B). In contrast to what has been reported for *E. coli*, *P. aeruginosa*, and *M. tuberculosis*, where maximal efficiency was observed upon targeting the non-template DNA strand close to the transcription start site, we observed that 4/5 guides, including the template-targeting GFP_g3F, performed similarly within each test condition (Figure 2B) [23–25]. By contrast, sgGFP_g4F, which targeted the template strand at a distance from the translation start site (TSS), exhibited zero silencing and mimicked the empty vector control regardless of nisin induction time (Figure 2B). In conclusion, maximal silencing efficiency is achieved with pre-sensitization, where all non-template targeted guides perform similarly regardless of the distance from the TSS or the GC content.

**Figure 2.**
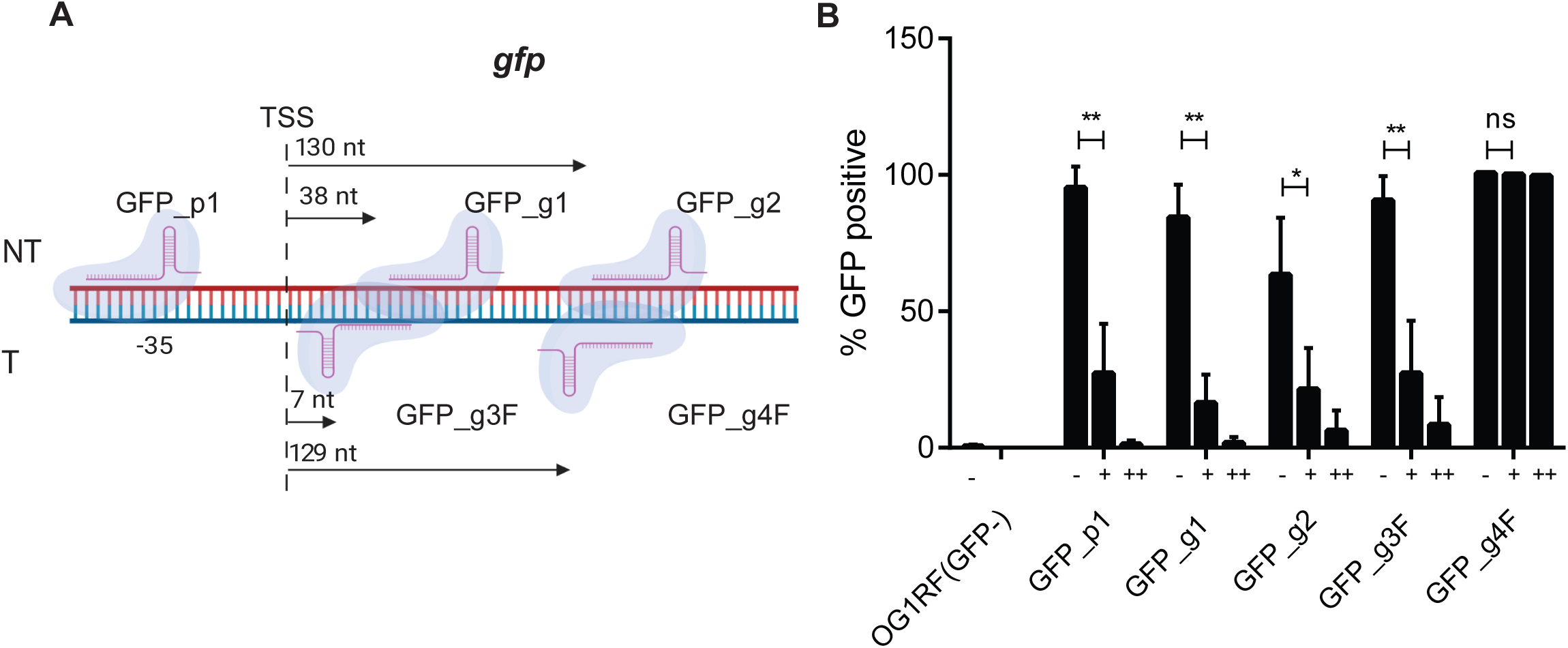
Efficient *gfp* silencing of pre-sensitized cultures on the non-template DNA strand. **A.** Schematic diagram of *gfp* operon indicates 5 sgRNAs that target the promoter region (GFP_p1), and protein-coding region on non-template (GFP_g1, GFP_g2) and template (GFP_g3F, GFP_g4F) DNA strands. Arrows indicate the distance from the translation start site to the first nucleotide of the bound gRNA. Image created with BioRender.com. **B.** The 5 sgRNAs were tested for *gfp* repression activity by pre-sensitizing bacteria with nisin overnight and sub-culturing with nisin induction the next day (++), or grown overnight without nisin and induced (+) or not-induced (-) the following day. After 2.5 hours of subculture, cells were washed and analysed by flow cytometry. The percentage of GFP expressing cells was determined by proprietary Attune NxT Flow Cytometer software from 500,000 events using EF_dCas9_ pp empty vector as a 100% positive control. Statistical analysis was performed by the unpaired t-test using GraphPad. **, P<0.001; *, P<0.05; ns, not significant.

### Efficient *ebpABC* operon and selective non-template DNA strand silencing in planktonic and biofilm bacteria

To explore the efficiency of CRISPRi silencing of whole operons, we designed sgRNAs to target the *ebpABC* operon, which encodes endocarditis and biofilm-associated pili (Ebp) important for biofilm formation [40, 41]. The *ebpABC* operon is comprised of *ebpA*, *ebpB* and *ebpC* expressed from the same promoter upstream of the *ebpA* [42]. EbpC is the major pilin subunit, EbpA is the tip adhesin, and EbpB is found at the base of the polymerized pilus [42]. In the absence of EbpA, long pili are still polymerized in which EbpC comprises the stalk of the pilus, while in the absence of EbpC only short EbpA-EbpB dimers are formed [43]. To test whether silencing the first gene in the operon could effectively silence the entire operon, we designed EbpA_g1 and EbpC_g1 that target the nontemplate protein-coding DNA strand 1845 and 5094 nt downstream of the *ebpA* TSS (Figure 3A). Since we observed selective efficiency of *gfp* template DNA strand targeting, to further explore this selectivity, we also designed EbpA_g2F and EbpC_g2F that target the template DNA strand 1157 and 6002 nt downstream the *ebpA* TSS, respectively (Figure 3A). We assessed the efficiency of operon transcriptional silencing by quantifying the amount of polymerized pili by Western blot and by quantifying pilus function in biofilm formation, since pilus deficient strains are attenuated for biofilm formation [40].

**Figure 3.**
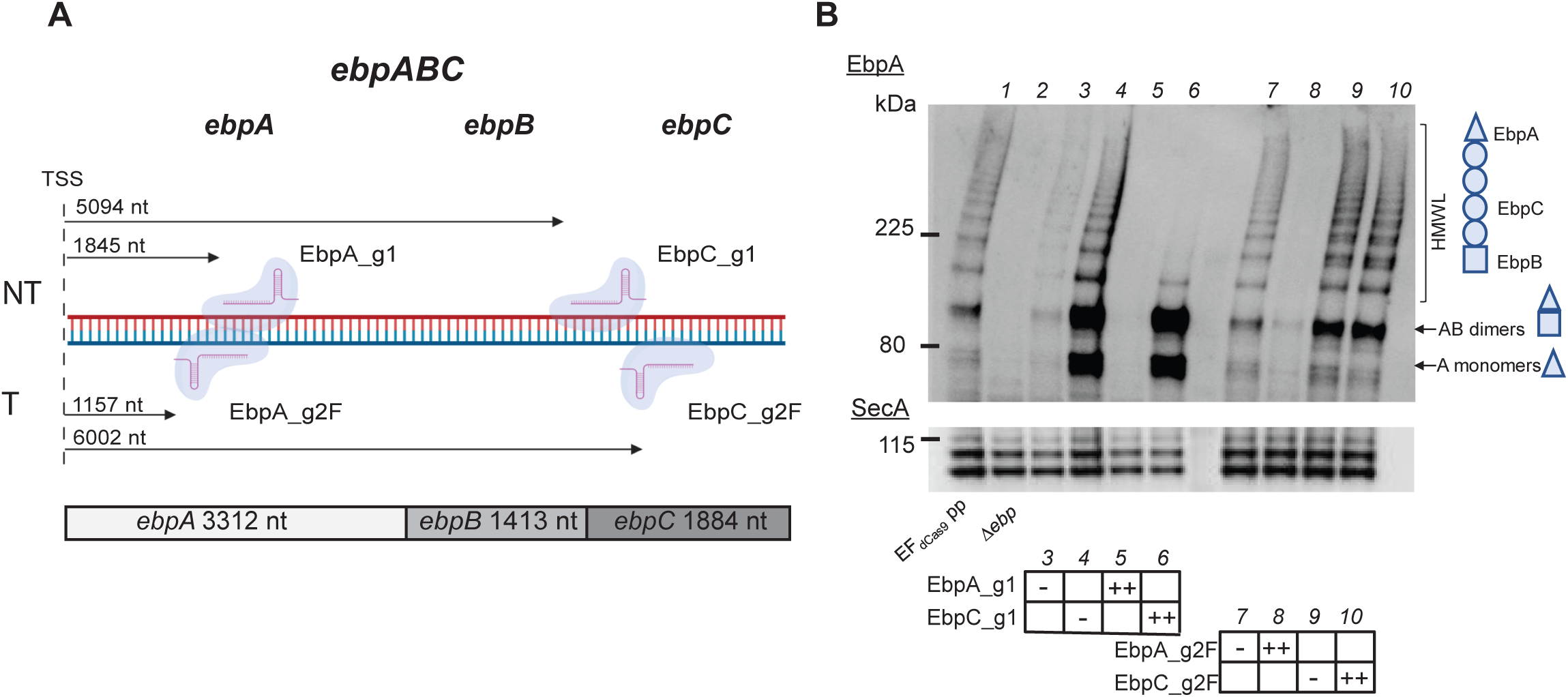
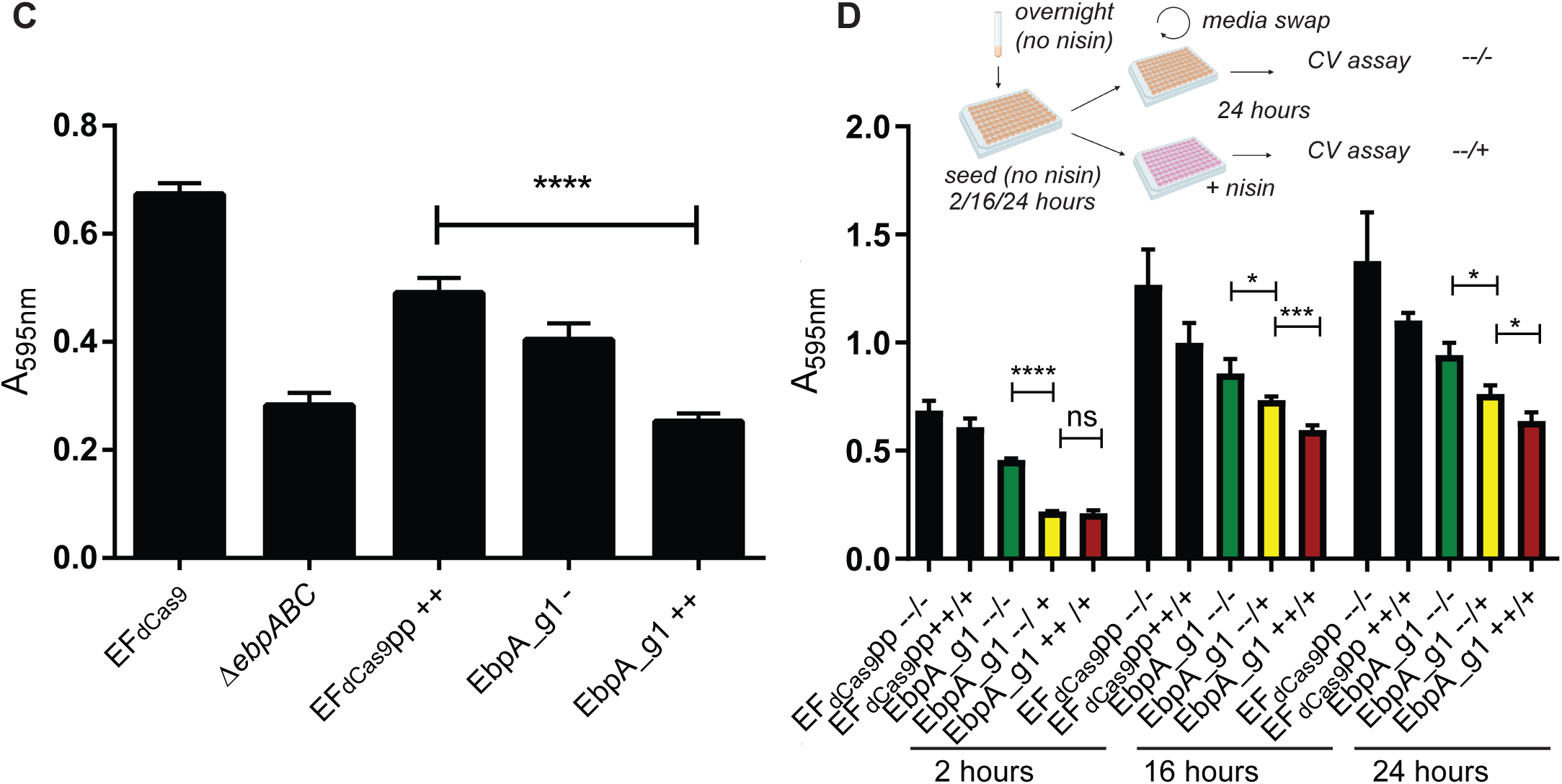
Efficient biofilm perturbation through CRISPRi targeting on *ebpABC*. **A.** Schematic diagram of *ebpABC* operon indicates 4 sgRNAs that target *ebpA* and *ebpC* protein-coding regions on non-template (EbpA_g1, EbpC_g2) and template (EbpA_g2F, EbpC_g2F) DNA strands. Arrows indicate the distance from the translation start site to the first nucleotide of the bound gRNA. **B.** Western blot probed with anti-EbpA antibody on whole cell lysates of *ebpA* and *ebpC* CRISPRi targeted strains, including ∆*ebpABC* and empty plasmid control strain (EF_dCas9_). Ebp appear as a high-molecular-weight ladder (HMWL) of covalently polymerized pili of different lengths. SecA served as a loading control and appears as double band ~100kDa **C.** Crystal violet staining of 24 hour biofilms formed on plastic in TSBG media. EF_dCas9_ pp and ∆*ebpABC* were used as controls, the test Ebp_g1 strain was uninduced (-) or induced with nisin (50 ng/ml) or pre-sensitized overnight and induced the following day (++) prior to seeding into biofilm chambers. Statistical analysis was performed by the unpaired t-test using GraphPad. ****, P<0.0001. **D.** Crystal violet staining on EbpA_g1 biofilms that were pre-grown on plastic without nisin induction for 2, 16 and 24 hours followed by media swap and 24 hours nisin induction (--/+, yellow bars). + and – indicate the presence or absence of nisin in the overnight culture, subculture and in the swap media. Swap is indicated as “/”. Constitutively induced cultures ++/+ (red bars) and constitutively uninduced cultures --/- (green bars) are the control strains. Statistical analysis was performed by the unpaired t-test using GraphPad. ****, P<0.0001; ***, P<0.001; *, P<0.05; ns, not significant.

Both guides that target *ebpA*, the first gene of the *ebpABC* operon, regardless of the targeted DNA strand, were similarly efficient in silencing the whole operon as measured by the absence of a high-molecular-weight ladder by Western blot (lanes 5 and 8, Figure 3A) indicating lack of pilus polymerization. As expected, lack of polymerized pili after EbpA_g1-mediated silencing correlated with reduced biofilm formation, similar to an *ebpABC* null mutant (Figure 3C). EbpC_g1 targeting the nontemplate strand efficiently silenced *ebpC* transcription, but did not affect expression of the upstream *ebpA* and *ebpB*, resulting in the absence of HMWL Ebp but presence of EbpAB dimers (lane 6, Figure 3B). By contrast, EbpC_g2F did not silence *ebpC*, leaving pili expression unaffected (lane 10, Figure 3B). Therefore, taken together with the *gfp* targeting data above, these data suggest that the silencing efficiency achieved by targeting the template DNA strand varies and, at least in these two instances, is less effective when the target is farther away from the TSS. By contrast, targeting the nontemplate strand is efficient to silence a single gene or the whole operon, and can act at a significant distance from the operon TSS.

Biofilm formation proceeds via a series of developmental steps, starting with adhesion of a single cell, aggregate or microcolony formation, maturation into heterogenous 3D structure, and ultimately dispersal [44]. Single gene knockouts or *a priori* gene silencing enable study of the contribution of gene products to biofilm adhesion or initiation; however, the study of specific genes to post-adhesion steps of biofilm formation has been more challenging to achieve. Since *ebpABC* is important in biofilm formation, we tested if CRISPRi can be used to perturb pre-formed biofilms to assess their involvement in *E. faecalis* biofilm maintenance. We allowed biofilms to form with uninduced bacteria for 2, 16 or 24 hours and subsequently swapped the media to fresh media with or without nisin and incubated the biofilms for another 24 hours. We observed a significant decrease in biofilm formation when nisin was added to silence pilus gene expression (--/+) when compared to uninduced biofilms (--/-) in early (2 hours) and matured (16 and 24 hours) biofilms (Figure 3D). Constitutively supressed Ebp (++/+) formed less biofilm as compared to post-biofilm formation induction (--/+)showing that Ebp are important for both initiation and maintenance of *E. faecalis* biofilm. These data demonstrate that inducible CRISPRi can be used to probe stage specific gene contributions to biofilm maturation and maintenance.

### CRISPRi silencing of *croR* mimics *croR*::Tn phenotype in antibiotic sensitivity assay

The CroRS two-component system contributes to *E. faecalis* antibiotic resistance, survival within macrophages, stress responses, and growth [45–49]. CroR phosphorylation by the cognate CroS sensor kinase is important for resistance to bacitracin, vancomycin and ceftriaxone, where ∆c*roR* cells are no longer resistant to these antibiotics [46]. To further validate the CRISPRi system in *E. faecalis*, we designed CroR_g1 to target *croR* on the non-template DNA strand and assessed sensitivity of the resulting strain to bacitracin, using *croR*::Tn as a control. Nisin-induced CroR_g1 bacteria mimicked *croR*::Tn and showed reduced resistance to bacitracin (MIC 8 µg/ml) as compared to the empty vector control (MIC 32 µg/ml) (Table 4). Hence, the CRISPRi system can be used to study genes involved in antibiotic resistance.

**Table 4.**
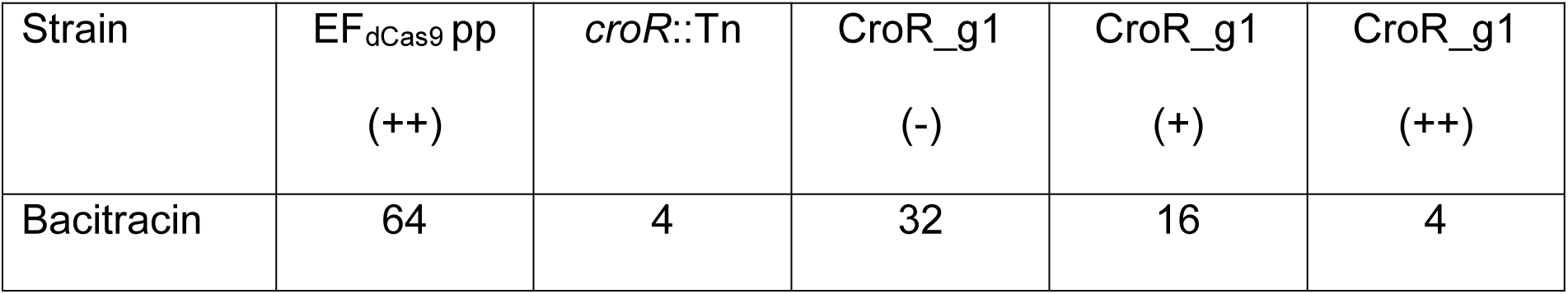
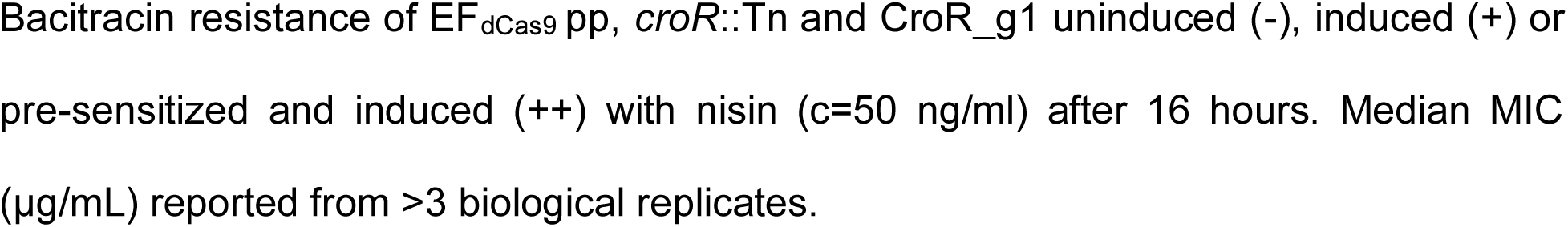
CRISPRi recapitulates antibiotic sensitivity phenotype of *croR* mutant.

| Strain | EF<sub>dCas9</sub> pp<br>(++) | *croR*::Tn | CroR_g1<br>(-) | CroR_g1<br>(+) | CroR_g1<br>(++) |
| --- | --- | --- | --- | --- | --- |
| Bacitracin | 64 | 4 | 32 | 16 | 4 |

Bacitracin resistance of EF<sub>dCas9</sub> pp, *croR*::Tn and CroR\_g1 uninduced (-), induced (+) or pre-sensitized and induced (++) with nisin (c=50 ng/ml) after 16 hours. Median MIC (µg/mL) reported from >3 biological replicates.

### Efficient combinatorial gene silencing

We next assessed the ability of the CRISPRi system to simultaneously silence multiple genes. At the same time we tested if the presence of a second guide with the same *nisA* promoter and Cas9 handle could be unstable or prone to recombination and interfere with the efficiency with CRISPRi as has been reported for some systems [50, 51]. We used combinatorial genetic *en masse* (CombiGEM) technology [28] to generate GFPEbpA_g1g1 and EbpASrtA_g1g1, enabling the expression of two sgRNAs under independent *nisA* promoters, to simultaneously silence *gfp* and *ebpA* or *ebpA* and *srtA* gene pairs. SrtA is an enzyme that covalently attaches polymerized pili to the cell wall, and deletion of this gene results in the absence of Ebp bound in the cell wall fraction of cells [42]. Upon nisin induction of GFPEbpA_g1g1, we observed the simultaneous loss of EbpA signal by Western blot (lane 8, Figure 4A) and reduction of GFP signal by flow cytometry (Figure 4B). Similarly, in the induced EbpASrtA_g1g1 strain, no signal was observed for EbpA or SrtA by Western blot (lane 6, Figure 4C), compared to the empty vector control (lane 1, Figure 4C). Thus, the combinatorial plasmids silenced two gene pairs with equivalent efficiency of a single-guide gene silencing, where the presence of a second sgRNA construct did not affect silencing efficiency of the other guide.

**Figure 4.**
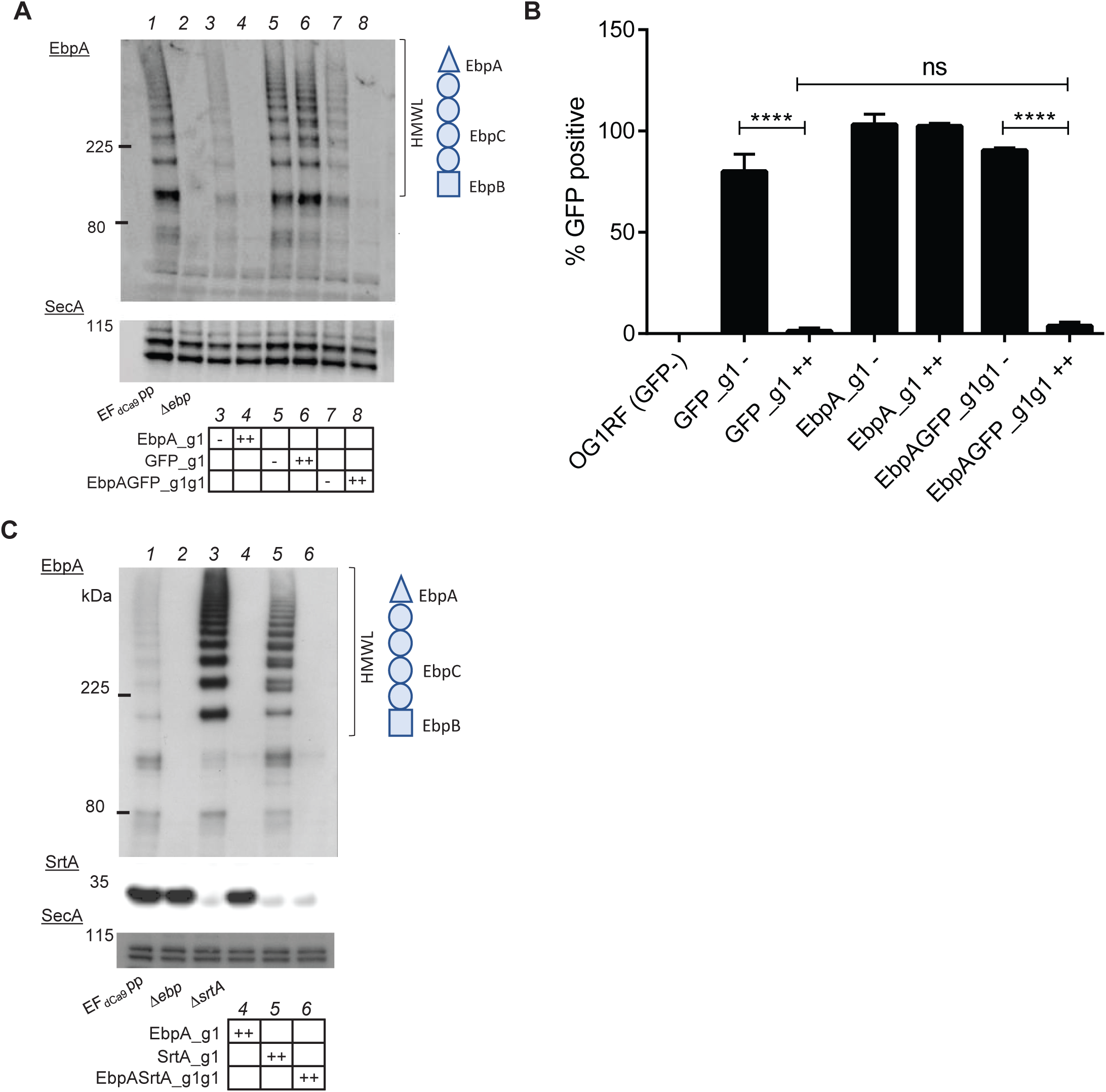
Efficient simultaneous silencing of two different genes. **A.** Western blot probed with anti-EbpA antibody on whole cell lysates of strains expressing EbpA_g1, GFP_g1 or EbpAGFP_g1g1 pre-sensitized and induced, *ebp* null and EF_dCas9_ pp were used as the control strains. SecA served as a loading control. **B.** Percentage of GFP-expressing cells as determined by proprietary Attune NxT Flow Cytometer software from 500 000 events from 3 independent experiments using EF_dCas9_ pp empty load as a 100% positive control. Statistical analysis was performed by the unpaired t-test using GraphPad. ****, P<0.0001; ns, not significant. **C.** Western blot probed with anti-EbpA and anti-SrtA antibodies on whole cell lysates of strains expressing EbpA_g1, SrtA_g1 or EbpASrtA_g1g1 pre-sensitized and induced cultures, EF_dCas9_ pp and ∆*srtA* were used as control strains. SecA served as a loading control.

### Essential gene targeting with pre-sensitizing at sub-inhibitory nisin concentration

Finally, we assessed the ability of the inducible CRISPRi to study essential genes. We chose to target *secA*, an essential gene of the general secretion pathway [52]. We designed SecA_g1 to target non-template DNA strand of *secA*. Because *secA* is essential, silencing of this gene is predicted to attenuate growth of the strain. We performed growth curves at various nisin concentrations to access growth inhibition after *secA* targeting. Even without induction we observed a reduced growth rate compared to the empty vector control, presumably due to a leaky *nisA* promoter with basal expression of dCas9_Str_ and SecA_g1 (Figure 5). Upon induction, we observed a similar growth inhibition for nisin concentrations 2.5-50 ng/ml, suggesting that maximal inhibition of SecA without pre-sensitization is achieved at 2.5 ng/ml (Figure 5B). To increase the degree of inhibition, we pre-sensitized bacteria overnight with 2.5 ng/ml nisin, as we observed no growth for overnight cultures grown at nisin concentration of 5-50 ng/ml (data not shown). When pre-sensitized cultures were further subcultured with 2.5 ng/ml of nisin we observed a more pronounced growth defect which was most apparent between 4-10 hours after induction, compared to the empty vector (no guide) control (Figures 5C). In summary, we showed that the function of essential genes can be studied using a low level nisin pre-sensitization and induction protocol.

**Figure 5.**
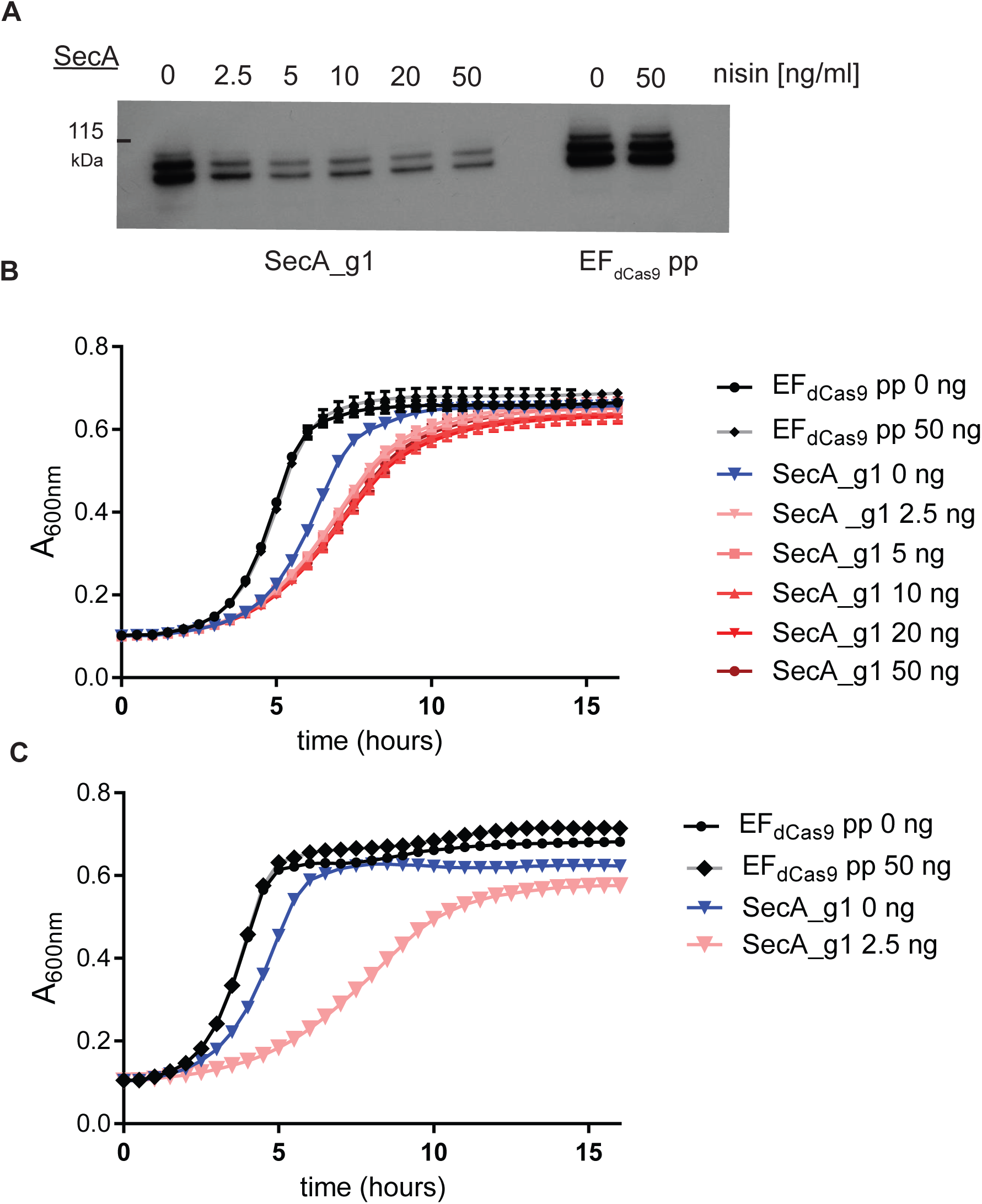
Essential gene targeting with pre-sensitizing at sub-inhibitory nisin concentration. **A.** Western blot probed with anti-SecA antibody on whole cell lysates from strain expressing SecA_g1 grown for 3 hours after subculture with nisin at concentrations 0-50 ng/ml. **B.** Growth curves of SecA_g1 induced at various nisin concentrations (0-50 ng/ml) without overnight pre-sensitization with EF_dCas9_ pp (empty load) as a control. **C.** Growth curves of SecA_g1 pre-sensitized overnight with 2.5 ng/ml of nisin prior subculture in a fresh media with nisin at 2.5 ng/ml. EF_dCas9_ pp (empty load) without the induction and with pre-sensitization and induction at 50 ng/ml were the controls.

## DISCUSSION

Genetic tools to easily and rapidly study the contribution of single or multiple enterococcal genes in a given biological process are lacking. To address this, we developed a scalable dual-vector nisin inducible CRISPRi system for *E. faecalis*. The system is most efficient on pre-sensitized cultures and can be used to study a variety of bacterial phenotypes, including biofilm formation, antimicrobial resistance, and gene essentiality.

Similar to CRISPRi systems developed for other bacterial species, our system is inducible, efficient, and can be multiplexed [21, 24]. We employed a two-plasmid system; one plasmid encodes dCas9 and a nisin responsive two-component system, and the other encodes the nisin-inducible sgRNA. The second plasmid, pGCP123, is small and easily modifiable for combinatorial targeting. We can introduce sgRNA into digested plasmid in the form of a gBlock (or reannealed oligos) through Gibson Assembly or In-Fusion reactions [22, 28, 53]. The plasmid can be further modified for simultaneous targeting of multiple genes by the ligation-digestion reaction of compatible restriction sites using CombiGEM technology [28]. Our system uses the streptococcal dCas9 with the handle, a well-characterized tool for CRISPRi [19, 23]. Although Cas9_Str_ is orthologous to native enterococcal Cas9 and shares the same PAM, we showed that the streptococcal dCas9 handle is not recognized by native enterococcal dCas9 and can be used in *E. faecalis* without modifying the endogenous Cas9 [26, 54].

We observed the highest dCas9_Str_ protein expression at 25 ng/ml of nisin. However, nisin is more stable at a lower pH and may partially degrade over the 24 hours in the culture media [55]. To account for the degradation, we used a higher nisin concentration of 50 ng/ml to maintain the maximal strength of *nisA* promoter [32, 55]. At 50 ng/ml of nisin, most gram-positive and gram-negative bacteria are able to replicate as the minimal inhibitory concentration is typically >1000 ng/ml, allowing our system to be used in the context of multi-species interactions [56]. Since the nisin-controlled gene expression system is functional in wide-range of Gram-positive bacteria including *Lactococcus*, *Lactobacillus, Leuconostoc, Streptococcus and Enterococcus*, therefore our CRISPRi-Cas9 can be potentially used in these genera [57].

To design sgRNAs, we used the CHOPCHOP database on the *E. faecalis* OG1RF genome and selected for the guides with a zero off-target score [29]. We tested sgRNAs of various GC-content (20-60%), targeting template and nontemplate DNA strands, and at various distances from the translation start site (TSS). The GC content played no role in the efficiency of silencing. All non-template DNA strand targeting sgRNAs were efficient, independent of the distance from the TSS, even though in *E. coli* the efficiency of the silencing was reported to decrease with increasing the distance from the transcription start site [19]. Surprisingly, we observed that 2 out of the 4 guides designed on the template DNA strand were still efficient in silencing the targeted gene, whereas it has been generally assumed CRISPRi sgRNA must target the non-template DNA strand for efficient silencing in bacteria [19–21, 23]. It is possible that template DNA strand-targeting guides, such as GFP_g3F, that bind to the template strand just 7 nt away from the TSS, may allow dCas9 to interfere with the assembly of the transcription machinery, preventing transcription initiation. Consistent with this possibility, another guide GFP_g4F that targets the template strand 200 nt downstream from the TSS does not silence GFP. However, targeting the template strand of *ebpA* using EbpA_g2F, which binds 1157 nt away from TSS, is efficient in gene silencing indicating that template strand targeting and silencing is not universally distance-dependent. Further work is needed to understand the nature and mechanism of template DNA strand silencing in *E. faecalis*.

A great challenge in studying gene contribution to different stages of developmental cycles, such as those that occur during biofilm formation, arises when early steps are essential for later steps to occur, necessitating the ability for stage-specific gene silencing. We leveraged the inducibility of our system to trigger *ebpA* silencing in pre-formed biofilms to address the role of these pili *after* biofilm initiation, during the maturation and maintenance stage. We observed significant reduction in biofilm biomass in the induced cultures compared to uninduced controls, indicating that CRISPRi/Cas9 can be used to perturb the pre-formed biofilms and to identify and interrogate gene targets in a biofilm stage-specific manner.

To expand the uses of this CRISPRi system, we utilized CombiGEM technology to generate, in one easy step, combinatorial plasmids to target two genes simultaneously [28]. We showed that the simultaneous expression of two guides was as efficient in silencing as the expression of a single guide per cell. Despite the presence of the same promoter sequence on different plasmids, we did not observe disruptive recombination of the promoters or between the guides, with stable and consistent repression of both targeted genes. Therefore, this system has the potential to be scaled up for sgRNA library preparation and high-throughput combinatorial studies.

Finally, our system was tested in the study of essential genes, where targeting the essential *secA* gene with minimal nisin induction significantly impaired bacterial growth, while high concentrations of nisin impeded the growth and killing the bacteria. Genetic tools to study essentials genes in *E. faecalis* are limited to transposon mutant library sequencing approach and essential gene inactivation with *in trans* complementation [6, 58, 59]. Therefore, we can deploy CRIPSRi to study gene essentiality under various nutrient conditions or leverage upon the systems inducibility and probe essentiality of the genes *in vivo*.

In summary, we have developed and validated an efficient CRISPRi system that can be readily used to study single or a combination of genes involved in different biological processes and can be modified for high-throughput screens, including combinatorial analyses, in *E. faecalis*. This tool will effectively facilitate the study of *E. faecalis* pathogenesis and allow rapid identification of novel targets for future interrogation.

## ACKNOWLEDGMENTS

This work was supported through core funding of Singapore-MIT Alliance for Research and Technology (SMART), Antimicrobial Resistance Interdisciplinary Research Group (AMR IRG). Part of the work was carried at Singapore Centre for Environmental and Life Science engineering (SCELSE) whose research is supported by the National Research Foundation Singapore, Ministry of Education to Nanyang Technological University and National University of Singapore, under its Research Centre of Excellence Programme. We thank Hooi Linn Loo and Peiying Ho (SMART AMR IRG) for assistance with flow cytometry. We thank Kline lab members Drs. Haris Antypas and Tom Watts for critical reading of the manuscript.

---

**Figure S1. gBlock sequence used to generate sgRNA expressing vector.**

The 20 nt sgRNA (N) transcribed from the *nisA* promoter (in green), followed by Cas9_Str_ scaffold sequence and transcription terminator (in red). Each guide was barcoded with a unique 6 nt barcode (B). The four restriction sites (underlined) were used for genetic assembly through CombiGEM. gBlock is flanked with overhang regions (in black) for InFusion reaction into PstI/KpnI digested pGCP123.

**Figure S2. Native enterococcal Cas9 does not interfere with nisin-inducible streptococcal dCas9**

**A.** Percentage of GFP-expressing cells from induced EF_dCas9_ Ebp_g2 with either empty pMSP3545 or pMSP3545-dCas9_Str_ determined by proprietary Attune NxT Flow Cytometer software from 500, 000 events from 3 independent experiments using EF_dCas9_ pp empty load as a 100% positive control. Statistical analysis was performed by the unpaired t-test using GraphPad. ****, P<0.0001.

**B.** Representative flow cytometry plots showing GFP expression in induced Ebp_g2 with pMSP3545 empty or pMSP3545-dCas9_Str_.

